# CETP inhibition and *ADCY9* genotype: evidence of a qualitative pharmacogenetic interaction in cardiovascular disease?

**DOI:** 10.1101/336875

**Authors:** Michael V Holmes, George Davey Smith

## Abstract

**Background:** CETP inhibitors raise circulating concentrations of HDL-cholesterol, and potent inhibitors also lower non-HDL-cholesterol and risk of vascular disease. Previous genome-wide pharmacogenetic analysis of a phase III randomized controlled trial (RCT) of the CETP inhibitor, dalcetrapib, found variants in *ADCY9* to associate with response to treatment. More recently, findings from a pharmacogenetic analysis of the CETP inhibitor evacetrapib reported a lack of such an association.

**Aims:** To clarify the totality of evidence on whether *ADCY9* genotype modifies the treatment response to CETP inhibition on risk of major adverse cardiac events through systematic review and meta-analysis.

**Methods:** We searched PubMed on 22^nd^ May 2018 to identify RCTs of CETP inhibition that reported vascular disease effect estimates stratified by *ADCY9* genotype. Stratum-specific estimates were pooled using fixed effect meta-analysis. Tests of heterogeneity between, and trend across, genotypic strata were assessed using Chi2.

**Results:** Nine studies were identified from PubMed, of which two (dal-OUTCOMES and ACCELERATE) were RCTs reporting the treatment response to CETP inhibition by *ADCY9* genotype, and fulfilled the inclusion criteria. In meta-analysis of dal-OUTCOMES and ACCELERATE, treatment with a CETP inhibitor was associated with a relative risk (RR) for major adverse cardiac events of RR 0.80 (95%CI, 0.65-0.99) in carriers of *ADCY9* rs1967309 AA. For carriers of AG, the corresponding estimate was a RR of 1.01 (95%CI, 0.89-1.13), and for GG carriers, it was RR 1.21 (95%CI, 1.06-1.40). We identified evidence of heterogeneity (P=0.004) and a trend (P=0.0009) across genotypic groups.

**Conclusions:** In contrast to the interpretation provided by authors of the analysis based in the ACCELERATE trial, the available evidence lends weak support to a potential interaction of CETP treatment by *ADCY9* genotype on risk of major adverse cardiac events. Additional data, e.g. from the ongoing dal-GenE trial focused explicitly on this interaction, should provide further clarity regarding the robustness of this pharmacogenetic effect.

---

Nissen and colleagues(1) conducted a pharmacogenetic analysis of a matched nested case control subset of the ACCELERATE trial of evacetrapib versus placebo to investigate whether a genetic variant in *ADCY9* (a single nucleotide polymorphism, SNP, rs1967309) modified treatment response. The study was motivated by a GWAS of dal-OUTCOMES, which identified *ADCY9* rs1967309 to associate with the response to another CETP inhibitor, dalcetrapib(2).

To clarify the pooled pharmacogenetic effect of *ADCY9* genotype on the treatment response to CETP inhibition versus placebo with risk of major adverse cardiac events, we performed a literature search and meta-analysis. We searched PubMed on 22^nd^ May 2018 using the search term “ADCY9” AND (“CETP inhibition” OR “CETP inhibitor” OR “evacetrapib” OR “anacetrapib” OR “dalcetrapib” OR “torcetrapib” OR “obicetrapib”). This identified 9 articles that we screened for inclusion. Criteria for inclusion were: (i) studies of humans; (ii) use of a randomized controlled trial design; (iii) intervention with a CETP inhibitor versus placebo; (iv) measurement of *ADCY9* genotype; and, (v) an assessment of whether *ADCY9* genotype modified the treatment response to CETP inhibition. This search identified the two trials (dal-OUTCOMES(2) and ACCELERATE(1)) and no further studies met the inclusion criteria. Using the reported data in the two trials, we conducted a fixed-effect meta-analysis of the treatment response to CETP inhibition stratified by *ADCY9* rs1967309 genotype. We tested for statistical evidence of heterogeneity in the treatment response to CETP inhibition using the Chi2 test for heterogeneity (2 degrees of freedom) and conducted a test for trend using the Chi2 test for trend (1 degree of freedom). Analyses were conducted in STATA V13.1 and R.

In meta-analysis of dal-OUTCOMES and ACCELERATE (Figure), treatment with a CETP inhibitor was associated with a relative risk (RR) for major adverse cardiac events of RR 0.80 (95%CI, 0.65-0.99) in carriers of *ADCY9* rs1967309 AA. For carriers of AG, the corresponding estimate was a RR of 1.01 (95%CI, 0.89-1.13), and for GG carriers, it was RR 1.21 (95%CI, 1.06-1.40). The test against the null hypothesis of no heterogeneity between *ADCY9* genotype groups was P=0.004 and P for trend was 0.0009.

**Figure.**
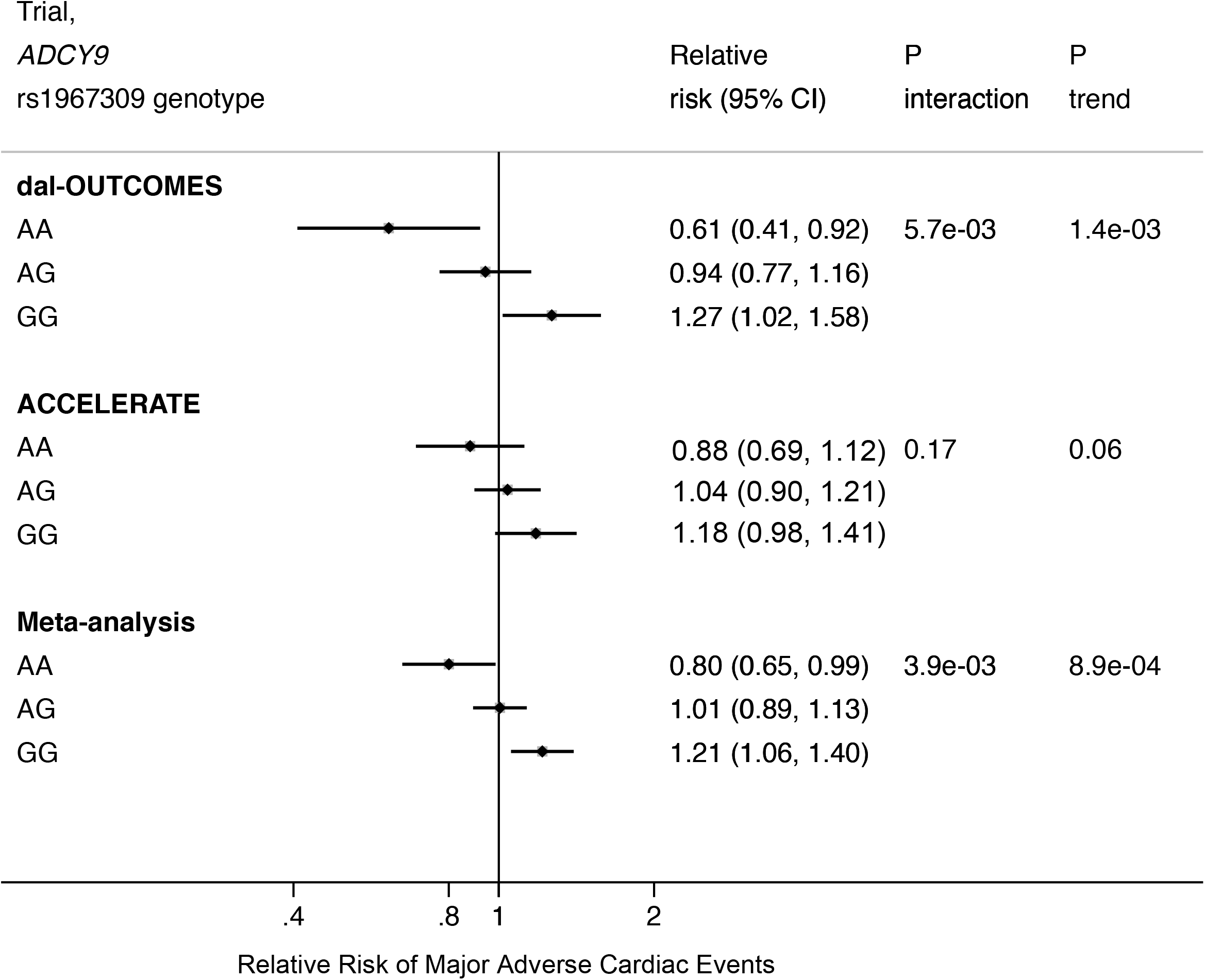
Fixed-effect meta-analysis of CETP treatment versus placebo and risk of major adverse cardiac events stratified by *ADCY9* rs1967309 genotype. P for interaction from Chi^2^ test for heterogeneity (2 degrees of freedom). P for trend from Chi^2^ test for trend (1 degree of freedom).

This meta-analysis provides some evidence of a potential qualitative pharmacogenetic interaction between CETP treatment and *ADCY9* genotype. In the study by Nissen et al(1), as the P-values for heterogeneity and trend were greater than 0.05, they concluded “Pharmacogenetic analysis did not show a significant association between the *ADCY9* SNP (rs1967309) and cardiovascular benefit or harm for the cholesteryl ester transfer protein inhibitor evacetrapib.” Such an appraisal, based solely on P-values doesn’t consider that the pharmacogenetic analysis in ACCELERATE was a replication test, and thus, effect modification of the treatment response to CETP inhibition by *ADCY9* rs1967309 genotype would not need to meet the same stringent thresholds as the initial discovery study. Furthermore, the tendency for the scientific community to dichotomize interpretations of findings into being positive or not, can lead to an overly simplistic interpretation. Following the publication of the study, media reports echoed this interpretation, e.g. “Precision Medicine Gamble Fails for CETP Inhibitors in

ACCELERATE Analysis” (https://wwwtctmdcom/news/precision-medicine-gamble-fails-cetp-inhibitors-accelerate-analysis). Our meta-analysis highlights the need to appraise the wider evidence base and suggests that, if anything, the findings by Nissen and colleagues could be cautiously interpreted as supportive, rather than as being “unable to replicate findings”.

In addition, Nissen and colleagues conducted analyses adjusted for cardiovascular risk factors. They note that the evidence for an interaction was attenuated on adjusting for conventional risk factors. While the *ADCY9* variant did not associate with risk factors in ACCELERATE or the UK Biobank, any attenuation on such an adjustment does not have an intrinsic meaning since alterations in effect could arise from adjusting for a mediator (which might induce collider bias).

In conclusion, the totality of available evidence lends weak support to a potential interaction of CETP treatment by *ADCY9* genotype. As the accompanying Editorial notes(3), additional data from pharmacogenetic analyses of the REVEAL trial(4), together with the dal-GenE trial(5) focused explicitly on this interaction, should provide further clarity regarding the robustness of this pharmacogenetic effect.

## References

1. Nissen SE, Pillai SG, Nicholls SJ et al. ADCY9 Genetic Variants and Cardiovascular Outcomes With Evacetrapib in Patients With High-Risk Vascular Disease: A Nested Case-Control Study. JAMA Cardiol 2018.

2. Tardif JC, Rheaume E, Lemieux Perreault LP et al. Pharmacogenomic determinants of the cardiovascular effects of dalcetrapib. Circ Cardiovasc Genet 2015;8:372–82.

3. Sabatine MS. Pharmacogenetics and the promise of personalized medicine. JAMA Cardiology 2018.

4. HPS3/TIMI55-REVEAL Collaborative Group, Bowman L, Hopewell JC et al. Effects of Anacetrapib in Patients with Atherosclerotic Vascular Disease. N Engl J Med 2017;377:1217–1227.

5. Tardif JC, Rhainds D, Rheaume E, Dube MP. CETP: Pharmacogenomics-Based Response to the CETP Inhibitor Dalcetrapib. Arterioscler Thromb Vasc Biol 2017;37:396–400.

